# Detecting host-parasitoid interactions in an invasive Lepidopteran using nested tagging DNA-metabarcoding

**DOI:** 10.1101/035071

**Authors:** James JN Kitson, Christoph Hahn, Richard J Sands, Nigel A Straw, Darren M Evans, David H Lunt

## Abstract

Determining the host-parasitoid interactions and parasitism rates for invasive species entering novel environments is an important first step in assessing potential routes for biocontrol and integrated pest management. Conventional insect rearing techniques followed by taxonomic identification are widely used to obtain such data, but this can be time consuming and prone to biases. Here we present a Next Generation Sequencing approach for use in ecological studies which allows for individual level metadata tracking of large numbers of invertebrate samples through the use of hierarchically organised molecular identification tags. We demonstrate its utility using a sample data set examining both species identity and levels of parasitism in late larval stages of the Oak Processionary Moth (*Thaumetopoea processionea* - Linn. 1758), an invasive species recently established in the UK. Overall we find that there are two main species exploiting the late larval stages of Oak Processionary Moth in the UK with the main parasitoid (*Carcelia iliaca*-Ratzeburg, 1840) parasitising 45.7% of caterpillars, while a rare secondary parasitoid (*Compsilura conccinata*-Meigen, 1824) was also detected in 0.4% of caterpillars. Using this approach on all life stages of the Oak Processionary Moth may demonstrate additional parasitoid diversity. We discuss the wider potential of nested tagging DNA-metabarcoding for constructing large, highly-resolved species interaction networks.

## Introduction

Invasive species are a growing global threat and pose a major risk to both natural and cultivated ecosystems with detrimental effects including direct competition for resources (Peck *et al.* 2014), predation on native species (Boland 2004) and even disruption of intentionally released biocontrol agents (Schooler *et al.* 2011). Economically, it is estimated that invasive species have a total global cost of at least US$ 70 billion annually (Bradshaw *et al.* 2016). In Europe, there are over 1590 non-native invasive arthropod species (estimate as of Roques 2010) and the rate at which species are establishing is increasing with ‘an average of 10.9 species per year for the period 1950–1974 to an estimated 19.6 species per year for 2000–2008’ (Roques 2010; Roques *et al.* 2016). The UK has the third largest non-native species burden in Europe, with more than 502 arthropod species and 1376 higher plants (Roy *et al.* 2014). Determining how to deal with invasive species is critically important for both ecological and financial reasons. Although good biosecurity is likely to be far cheaper than control or eradication of established species (Vazquez-Prokopec *et al.* 2010), this is not always achieved. Thus, an understanding of the interactions between invasive species and native species within an invaded range is essential for quantifying the impacts on communities (Roy *et al.* 2009; Hesketh *et al.* 2010) as well as developing practical management approaches, such as biocontrol. Ecological network modelling provides a framework within which these questions can be addressed but, typically requires well sampled networks that are laborious to create with traditional observational approaches (Evans *et al.* 2016). Molecular tools and in particular, modern sequencing technologies, provide a way to collect this data on a large scale by standardising and automating much of the effort required to detect interactions (Handley *et al.* 2011).

The Oak Processionary Moth (*Thaumetopoea processionea*, hereafter referred to as OPM) is historically considered a native of the warmer parts of southern and central Europe with a more sporadic presence in western Europe (Groenen & Meurisse 2012) but since the 1970s the frequency of outbreaks in north-western Europe (especially Belgium, the Netherlands and Germany) has increased dramatically with multiple intense outbreaks where the gregarious larvae reach population densities of thousands of individuals per host tree (deciduous *Quercus spp.*) (Stigter *et al.* 1997). Population densities on this scale are capable of defoliating large areas of oak forest (Wagenhoff & Veit 2011). In addition to commercial forestry concerns, the caterpillars are also a serious public health risk due to the presence of urticating hairs containing thaumetopoein, a strong allergen unique to OPM and related moth species (Lamy *et al.* 1986).

The Oak Processionary Moth arrived in the UK in 2006 and has spread throughout Greater London but has yet to establish beyond this area (Mindlin *et al.* 2012). This is in part due to a UK Government control program that involves both manual nest removal and insecticide spraying using *Bacillus thuringiensis* (*Bt*) and even though the government funded program does not cover the complete UK range of the moth, the costs are still high at around £1.2 million in 2016 (Forest Research 2017). With costs this high, it is desirable to find an alternative to the current control method that is both financially viable and minimises any adverse ecological impact. Natural enemies of OPM have been investigated in Europe and at least 30 egg, larval or pupal parasitoids are known (Zwakhals 2005; Sobczyk 2014; Roques 2014; Sands *et al.* 2015), but often these records are collected on an *ad hoc* basis and other than one incidence of a single parasitoid being collected in London [Richmond Park: *Carcelia iliaca (Sands et al. 2015)]*, nothing is known about parasitoids of OPM and their infection rates in the UK. Understanding which parasitoids utilise OPM and the parasitism rates for each parasitoid is essential for assessing the potential use of these species for Integrated Pest Management (IPM). Species interactions have traditionally been detected using either observational data (e.g. Maglianesi *et al.* 2014), microscopic analysis of collected specimens (e.g. gut contents Otte & Joern 1976; Hyslop 1980; or pollen on pollinators Lopezaraiza–Mikel *et al.* 2007) or the rearing of organisms to identify plant-herbivore and host-parasitoid interactions (Pocock *et al.* 2012), but these approaches are labour intensive and there are often significant taxonomic hurdles to overcome, both in terms of the knowledge base required to accurately perform identifications and the presence of cryptic species (e.g. Smith *et al.* 2007, 2008; Kaartinen *et al.* 2010). In addition to this, studying the parasitoids of OPM and other species with urticating hairs can make laboratory rearing impractical and there is evidence that more traditional rearing methods can underestimate parasitism rates (Day 1994). Thus a better method is required to understand host-parasitoid interactions, the vital first step in assessing the potential for biocontrol methods.

The advent of molecular biological tools has allowed unprecedented opportunities to determine hitherto difficult to observe species interactions. Most of the work to date has focussed on studies using PCR diagnostic approaches where a primer pair specific to a single species is used to amplify only that species within a more complex DNA mixture and then visualise a band on an agarose gel. This is typically applied to known sets of species by using suites of primer pairs that each produce bands of different lengths that can then be separated by gel or capillary electrophoresis (e.g. aphid - parasitoid interactions Traugott *et al.* 2008). These approaches are extremely targeted and require extensive *a priori* knowledge regarding the interacting species as specific primers must be designed for each target species.

Massively parallel ‘next generation’ sequencing (NGS) is now a commonly used tool in diverse areas of ecology and has the advantage of being able to separate mixtures of DNA from multiple species into their constituent components. One commonly applied approach is ‘community metabarcoding’ where a bulk DNA sample from one environment is PCR amplified for a standard barcode locus, sequenced, and taxa comprising the community identified bioinformatically (e.g. Taberlet *et al.* 2012; Yu *et al.* 2012). Bulk sampling and sequencing a few complex samples results in community rather than individual level data. While this is useful for the detection of species (e.g. Dejean *et al.* 2012) or characterisation of whole communities across treatments or time (e.g. Yu *et al.* 2012; Giguet-Covex *et al.* 2014), this is not necessarily appropriate for detecting species interactions (but see Leray *et al.* 2013 for a dietary analysis using this approach). The ideal species interaction detection method would involve the ability to sequence a wide variety of organisms in complex mixtures (e.g. extracted DNA containing both host and parasitoid) while retaining individual sample level metadata so that semi and fully quantitative networks can be created *sensu* Hrček and Godfray (2015).

The use of unique MID tags (Molecular Identification tags, 8-mer oligonucleotide sequences) added to the PCR primers is a well-tested strategy for sample tracking in multiplexed samples with NGS approaches (Binladen *et al.* 2007). Eight forward MID tags are typically matched with twelve reverse MID tags to give 96 unique tag combinations. Multiple sets of primers like this can increase the number of samples used but large numbers of unique MID-labelled primers can be expensive and complex to organise in a laboratory environment, making it unusual to have more than four sets of primer combinations in a single experiment (384 samples although see Campbell *et al.* (2015) for an example of highly multiplexed SNP genotyping). Sequencing 384 individual insects per sequencing run using a next generation sequencing approach to detect species interactions is possible, but due to the cost per sample it is unlikely to be feasible for the thousands of samples required when building well sampled ecological networks (Evans *et al.* 2016).

Shokralla *et al.* (2015) and Cruaud *et al.* (2017) provided a potential solution to this problem by utilising a nested barcoding approach involving two PCR steps followed by sequencing on the Illumina MiSeq that allow for the tracking of a large number of individual samples. However, Shokralla *et al.* provided limited reproducibility for readers with their workflow and Cruaud *et al.* focussed on mass barcoding of samples for species delimitation and construction of identification databases. To date, no studies have demonstrated the utility of these approaches for detecting species interactions, although see Lefort *et al.* (2017) and Šigut *et al.* (2017) for examples of parasitism using more conventional sample tracking methods

In this study we have two interlinked objectives: 1. To present a simplified version of nested-metabarcoding methods for determining OPM-parasitoid interactions using a single PCR locus for a large number of samples, including improvements to control cross contamination; and 2. Demonstrate the utility of nested metabarcoding for detecting species interactions by determining the parasitoid identities and rates for one recently established population of OPM in the UK using a reproducible pipeline for creating host-parasitoid networks (metaBEAT 0.97.7 https://github.com/HullUni-bioinformatics/metaBEAT) with a downloadable working environment conveniently packaged in Docker (Docker Inc. 2017) and GitHub (GitHub Inc. 2017). By combining these objectives, we discuss how the ability to link metadata to individuals opens many new avenues for research including the ability to create larger more highly-resolved ecological networks for habitat management and restoration (Evans *et al.* 2016).

## Methods

### The nested metabarcoding approach

We employed a modification to the standard Illumina 16S bacterial metabarcoding protocol (Illumina 2011). In the original protocol two rounds of PCR were used to: firstly to isolate, and amplify the gene region of interest (PCR1) and; secondly, to add a set of molecular identification tags (MID tags) and the Illumina MiSeq adapter sequences (PCR2). Our modifications to the protocol include adding (1) an additional set of MIDs in PCR1 to further increase the resolution of sample identification, and (2) adding a sequencing heterogeneity spacer to improve MiSeq performance (Fadrosh *et al.* 2014). Each MID tag was composed of a unique 8-nucleotide sequence allowing them to be bioinformatically linked back to the individual sample. We included MIDs in both the forward and reverse primers with twelve forward tags and eight reverse tags, to give 96 unique combinations of sample tags that can be arranged on a single plate (See Fig. 1 for general primer design). We also included specific MID tags for positives and negatives as this helped track contamination and mistagging through illegal tag combinations. A plate of PCRs with these tagged primers was carried out with each PCR well being given a unique combination of tags. The PCR products were then pooled into a separate pre-library for each plate of samples (PCR1, Fig2). The pre-library was then used as a template for a second round of PCR which added the adapters necessary for Illumina sequencing. This reaction also added two additional MID tags that uniquely identify the plate (PCR2, Fig2). These tagged pre-libraries could then be purified, pooled and sequenced on a single Illumina MiSeq run.

**Figure 1.**
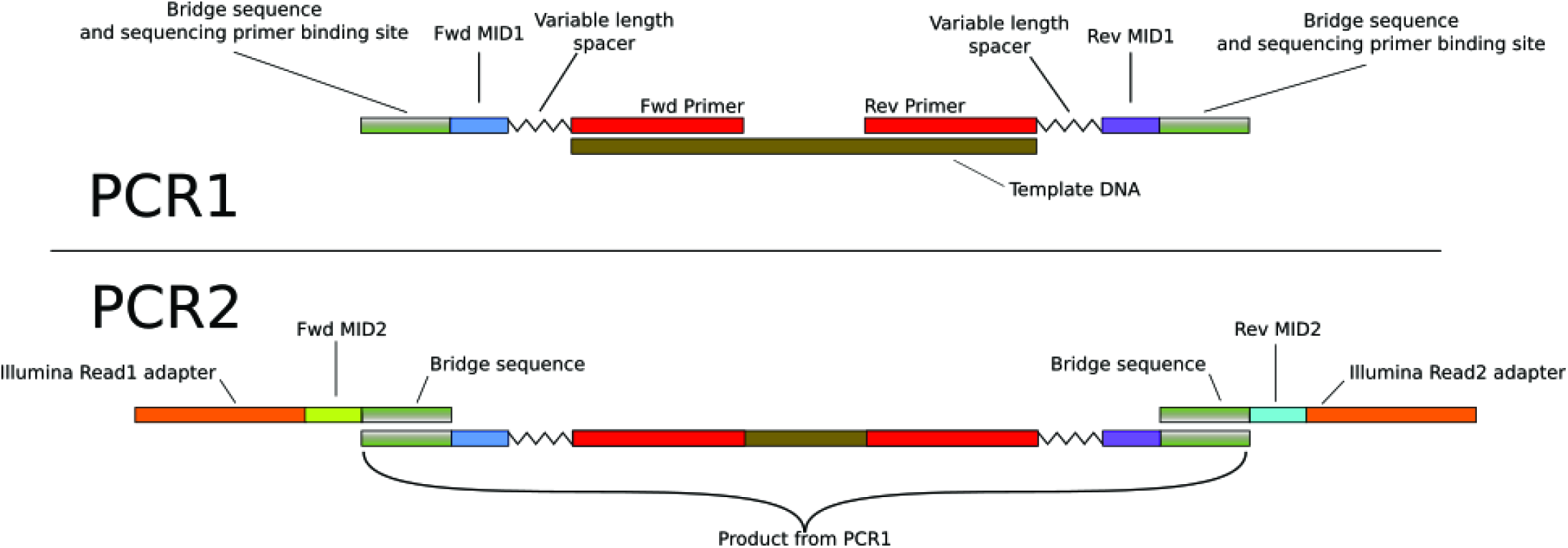
A simplified primer structure for a nested metabarcoding approach to detecting species interactions.

### Sampling and laboratory protocols

For this study 1012 OPM caterpillars (4th to 6th instar) were extracted from 26 nests (silk structures created by communally living caterpillars) collected from various locations in Croydon, London, UK in July 2014 (full collection data is available in <u>Table S1 in the Github repository</u>). Nests were frozen whole at −20°C for at least 48 hours to kill the caterpillars before the nest was opened up and individual caterpillars removed. Whole caterpillars were placed in deep well plates with a single 5mm stainless steel ball bearing per well and 300 μl of digestion buffer one (20mM EDTA, 120mM NaCl and 50mM Tris). Mechanical lysis was then performed by shaking in a Qiagen TissueLyser II for 2 x 2 minutes at 30Hz. The caterpillar slurry was centrifuged to remove tissue residue from lids and reduce the possibility of cross contamination. To each sample, 270 μl of digestion buffer two (20mM EDTA, 120mM NaCl, 50mM Tris and 2% SDS) plus 30 μl of 10 mg/ml Proteinase K solution were added. The plates were then mixed by repeated inversion and digested overnight at 37^o^C. After enzymatic lysis, 10 μl of the digestion supernatant was then used as the starting material for a 70 μl HotSHOT DNA extraction (Truett *et al.* 2000) which was diluted 1/100 for PCR amplification.

A 313 bp fragment of the Cytochrome C Oxidase subunit I barcode region (*coxI*) was amplified using primers based on mICOIintF and jgHCO2198 modified from Leray (2013) to include standard Illumina MIDs and bridge sequences (see Fig. 1 and <u>Table S2 in the GitHub repository associated with this manuscript</u>) along with a variable length sequencing heterogeneity spacer as in Fadrosh (2014). PCRs were carried out over 45 cycles (95°C for 15s, 51°C for 15s and 72°C for 30s) in 20 μl reactions using a high fidelity *Taq* mastermix (MyFi Mix Bioline), 1 μl of template DNA and each primer (final concentration - 0.5 μM). Extra cycles were required as long primers are known to cause a lag in PCR amplification (Schnell *et al.* 2015). In order to prevent cross contamination between wells, all PCRs were performed in individually capped PCR strips and all wells were sealed using mineral oil. In addition to this, oil was placed in the PCR well before all other reagents and the PCR master mix was mixed with primers and template DNA under oil to prevent cross contamination. An example output from a poorer performing run not employing these methods can be seen in Supplementary information appendix 1.

PCRs were checked on a gel to gauge success rates and 10 μl of each product from a plate was pooled together (without quantification) to produce each pre-library, resulting in eleven separate pre-libraries. Two aliquots of each pre-library were gel purified to remove remaining primers using QIAquick gel extraction kit (Qiagen) while maximising library recovery. The second library PCRs were carried out over 12 two-step cycles (98oC for 20s then 72oC for 30s) in 20 μl reactions using MyFi Mix (Bioline), 5 μl of pre-library and each primer (final concentration - 1.0 μM). PCR cycles were minimised so that nested-tagged PCR products were formed with minimal additional PCR error. Identical contamination control procedures were employed for PCR2 as in PCR1 (Fig. 2). This resulted in eleven libraries each with a unique set of library MIDs and a set of sample MIDS repeated across libraries. Libraries were gel purified and concentrations were quantified on a Qubit 3.0 using the Invitrogen dsDNA HS Assay Kit before being pooled at equal concentrations. The final set of pooled libraries was denatured and loaded onto a MiSeq using a v2 (2x250bp) sequencing kit with a final concentration of 12 pM and 10% PhiX as a sequencing control.

**Figure 2.**
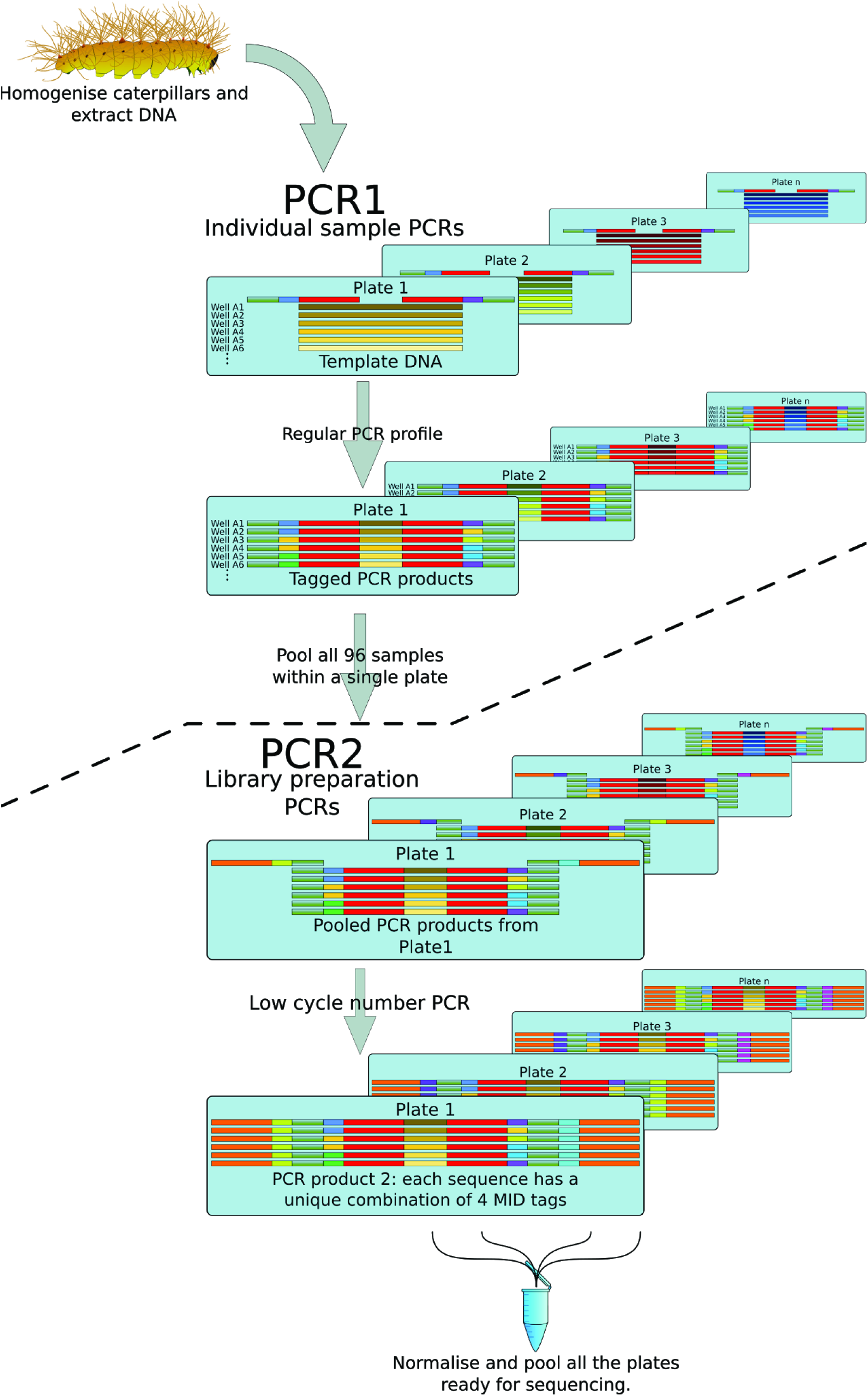
Preparation of PCR amplicon libraries for Illumina MiSeq using a nested metabarcoding approach to detecting species interactions. Colour choices have the same meaning as in Figure 1. Shades of colours represent the same target sequences in different individuals. Adapted from Evans *et al*. (2016).

Each plate contained 92 OPM samples, two negative samples (one HotSHOT extraction negative and one PCR water negative), and two positive samples. The first positive contained extracted template DNA from *Astatotilapia calliptera* (a cichlid fish) and was amplified at the same time as the OPM samples (hereafter denoted DNA positive). The second positive (hereafter denoted PCR positive) consisted of PCR products from *Mytilus edulis* (common mussel). Both positive samples were chosen due to their low probability of occurring in UK oak trees. The PCR positive was amplified independently from all other samples using primers with the correct combination of tags and was quantified using a Qubit 3.0 before being added directly to the pre-library during pooling. The PCR positive volume added to each pre-library was calculated so that we were adding 1/94th the total DNA (96 samples minus two negatives) of each pre-library as PCR positive. All samples were sequenced (including positives and negatives) even when no band was present as PCR products may still exist below gel detectable levels.

### Bioinformatic processing of Illumina MiSeq output

Processing of Illumina data from raw sequences to taxonomic assignment was performed using a custom pipeline for reproducible analysis of metabarcoding data metaBEAT v0.97.7. Individual steps performed as part of the pipeline are as follows: In brief, reads were demultiplexed using the *process_shortreads* script from the Stacks software suite (Catchen et al. 2013). Trimmomatic 0.32 (Bolger et al. 2014) was subsequently used for quality trimming and PCR-primer clipping of the raw reads in two steps: (1) reads were end-trimmed to phred Q30 using a sliding window approach (5bp window size) and (2) PCR-primers were conservatively clipped off the reverse complement sequences by removing 40bp from each read. Reads shorter than 100bp after quality trimming/primer clipping were discarded. Paired-end sequences were subsequently merged (minimum overlap 10bp) using FLASH 1.2.11 (Magoc & Salzberg 2011). Successfully merged reads were length filtered to retain only amplicons of the expected length (313bp +- 10%). The remaining high-quality sequences were reduced to unique sequences using vsearch v.1.1 (Rognes *et al.* 2016) by clustering with 100% similarity. Clustering results were further filtered based on the number of reads assigned to each cluster (minimum cluster coverage - see below for details) in order to minimize cross-contamination effects between wells. Surviving clusters in each well were then further clustered globally (again at 100% similarity) to reduce the number of BLAST searches performed. Single representative sequences from each cluster were subjected to a BLAST search (Zhang et al. 2000) against a local copy of the NCBI’s nucleotide database (nt). Sequences with at least 95 % similarity across at least 90% of their length to any sequence in the database were subjected to taxonomic assignment using a lowest common ancestor (LCA) approach similar to the strategy used by MEGAN (Huson et al. 2007), such that for each query we identified the taxa receiving the top 10% (bit-score) BLAST hits and subsequently determined the lowest taxonomic level shared by all taxa in the list.

### Assignment of thresholds for data processing

We employed three approaches to filtering the dataset and chose the most conservative approach for the final analysis. First, we examined our negative wells to see what minimum cluster coverage would effectively exclude background contamination (i.e. what is the minimum cluster coverage that results in zero retained reads and clusters in negative wells after filtering). Second, we examined our sample wells to look at what minimum cluster coverage resulted in stable per well read depths. We interpret this as the removal of minor components of each well such as PCR errors, mistagging errors and possible background contamination. Finally, we explicitly examined all possible illegal PCR1 MID tag combinations (four unused sample tag combinations plus a further 92 tag combinations involving special tags for positives/negatives and sample tags in all forward/reverse combinations). This allowed us to assess the minimum cluster coverage necessary to exclude sequences due to mistagging (i.e. possible tag swapping during PCR2 or signal bleed at the sequencing stage). For each approach we performed the clustering analysis across a range of minimum cluster sizes from 6 reads to 101 reads and plotted boxplots of both per well read depth and per well cluster retention (except for mistagging where we only examined per well read depth).

## Results

### PCR and sequencing success rates

Overall we had an apparent PCR success rate of 96.3% (i.e. 96.3% of sample wells produced a visible band on a gel). Analysis of clustering thresholds revealed that no negatives contained detectable reads or clusters above approximately 36 reads (Figs. S7A and S8A in appendix 3), and that per well read depth and clusters retained in the sample wells became stable at approximately 56 reads (Figs. S7B and S8B in appendix 3). The mistagging analysis revealed that the single largest cluster created by any illegal tag combination was 66 reads (Fig. S9 in appendix 3). Based on these results, we believe that performing our final analysis using a conservative minimum cluster size of 67 reads resulted in the effective exclusion of errors from the dataset.

From a single MiSeq v2 Illumina run we produced a total of 12,112,538 untrimmed sequences and retained 11,078,131 after quality trimming (91.5%). For the 1,012 moth samples, read depth per well ranged from 0 - 63,182 reads before quality trimming (mean = 11,840.1, sd = 9,309.6) and from 0 - 57,336 reads after quality trimming (mean = 10,870.9, sd = 8,613.8) (Fig. 3). Overall we had a sequencing success of 94.5% (percentage of sample wells for which reads were retained after data processing), eight out of eleven DNA positives sequenced successfully, but none of the PCR positives were successful. For the failed DNA positives, one produced clusters that were just below our minimum cluster coverage so was excluded, while the remaining two produced no reads at all. The PCR positives produced raw reads, but they were of poor quality leading to very few reads merging and all clusters being excluded (Fig. 4). The DNA positives that failed started with low raw read counts despite having distinct bands on agarose gels. This suggests that a pooling error led to underrepresentation of these wells in the final pooled pre-library which probably resulted in dropout. The PCR positives also produced strong consistent bands on a gel prior to being added to the pre-libraries, but because we quantified how much PCR positive we should add, we are less inclined to believe that a simple pooling error was responsible. A primer synthesis error in either the bridge sequence or the sequencing primer binding site for one or both of the PCR positive primers would result in complete sequencing failure. A sequence error in the bridge sequence would result in poor library formation for this sample during PCR2 while an error in the sequencing primer site would result in unreliable sequencing signal and subsequent filtering. To try and resolve this, Sanger sequencing of the PCR1 PCR positive product was undertaken. Reverse sequencing revealed that the forward primer was identical to the designed sequence. Forward sequencing was less successful and could not produce a strong Sanger trace (probably due to the primer length being suboptimal for Sanger sequencing). Alignments of the poor quality sequence and the reverse primer suggest there may be two deletions within the bridge sequence leading to poor library formation in PCR2. Further studies conducted subsequently were performed using newly synthesised PCR positive primers and sequencing was successful. We also recommend that primers for future studies should be synthesised at the highest possible quality standard to ensure accuracy of synthesis.

**Figure 3.**
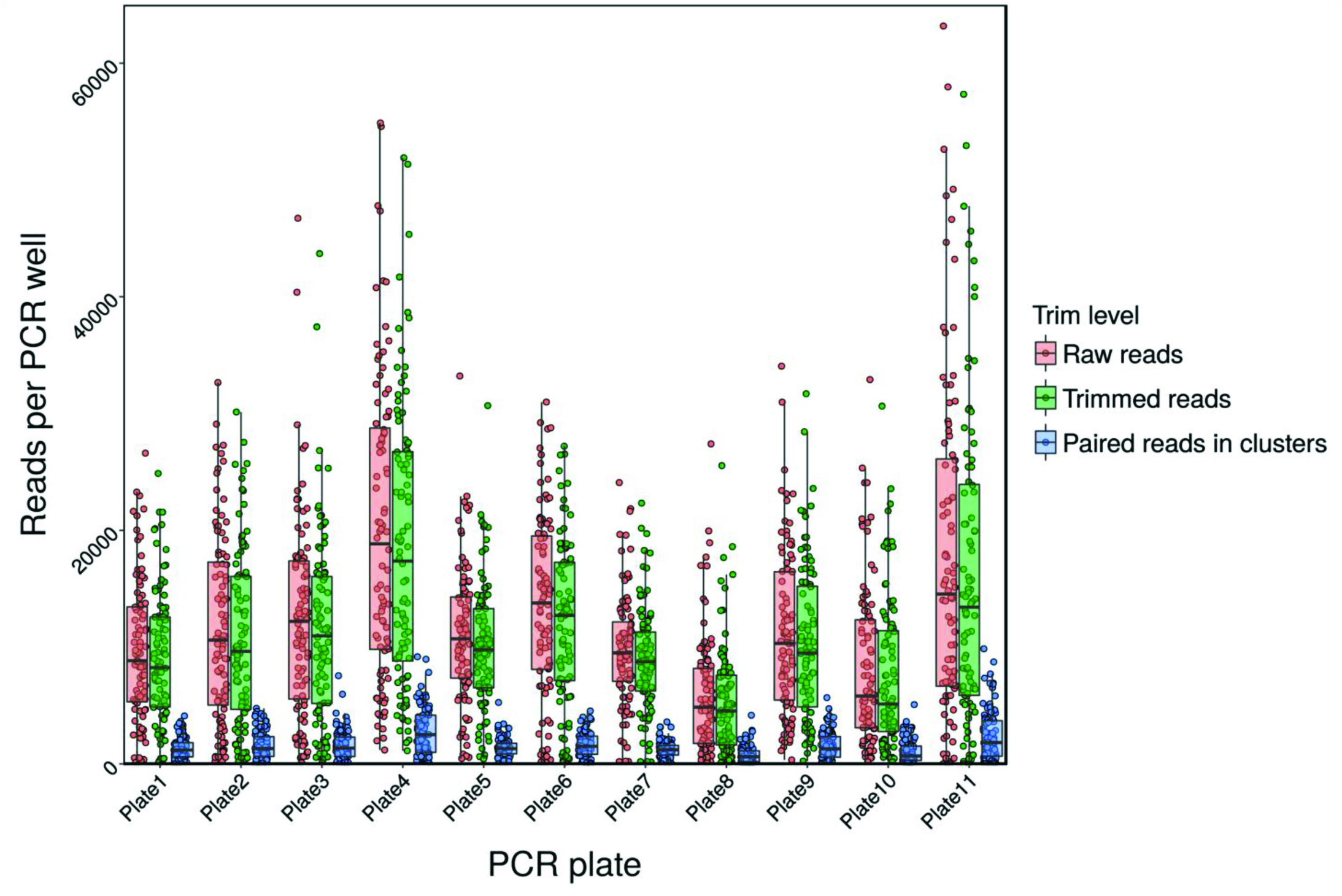
Read depth per PCR well for each plate (positives and negatives excluded) with actual read depth for each PCR well overlaid as scatter plots.

**Figure 4.**
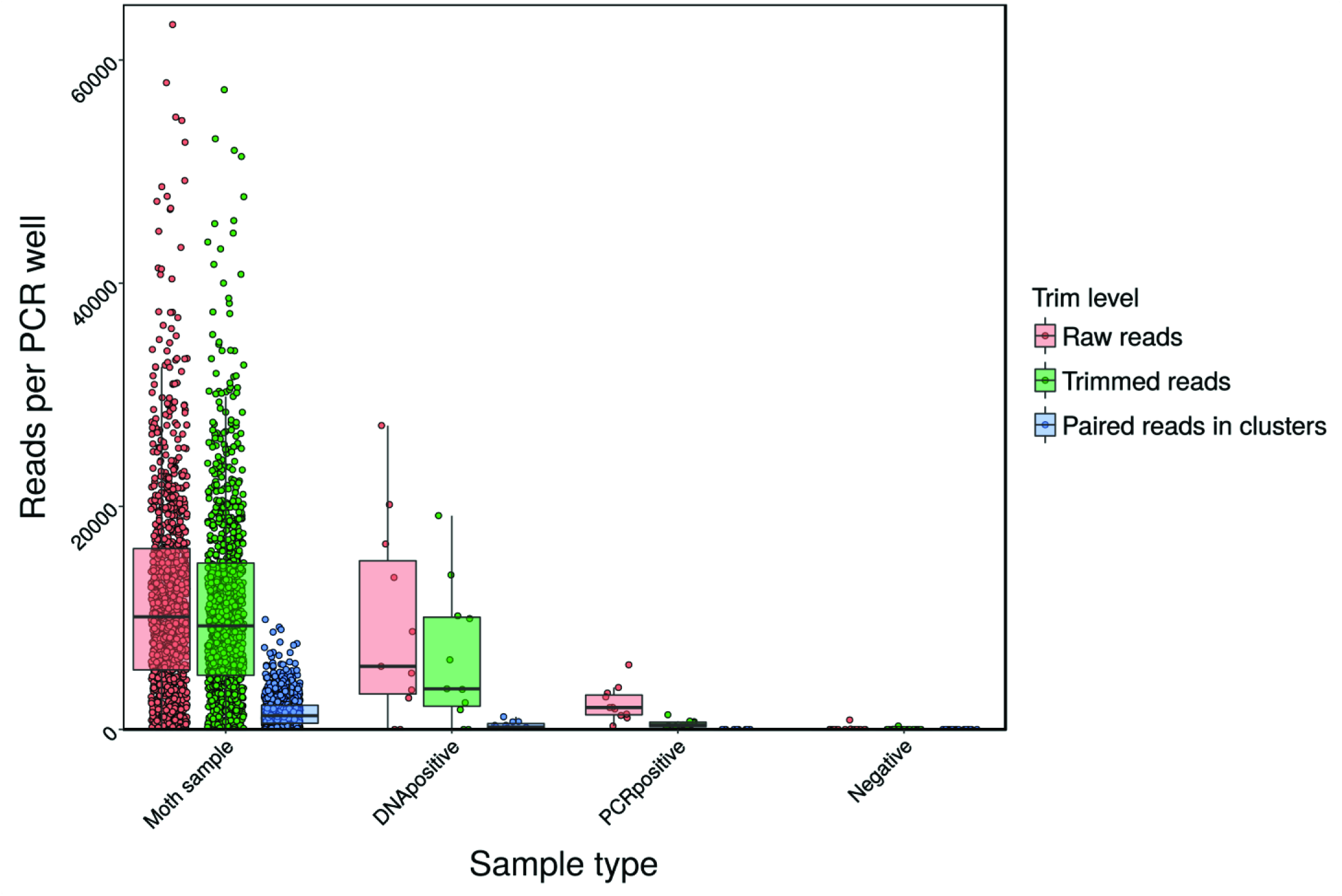
Boxplots of read depth per PCR well for each type of PCR well with actual read depth for each PCR overlaid as scatter plots.

### Species identifications and parasitism levels in OPM

Most sequences were assigned by BLAST to either OPM or its known parasitoid fly *Carcelia iliaca* (Sands *et al.* 2015) with an additional rare parasitoid *Compsilura concinnata* (a parasitoid normally associated with Gypsy moth -*Lymantria dispar* - Fig. 5A). Our data indicated that 45.7% of OPM caterpillars sampled from Croydon, London, were parasitised by *C. iliaca* while just 0.4% were parasitised by *Compsilura concinnata*. No Hymenopteran parasitoids were detected, so to check for non-amplification of Hymenoptera parasitoids by Leray primers we attempted PCR amplification of known pupal parasitoids of OPM and achieved 100% success (see Fig S6 in appendix 2 for further details), leading us to conclude that the Hymenopteran parasitoids tested were not present in this life stage of OPM. In addition to insect parasitoids we also detected a number of fungal sequences including the entomopathogenic ascomycete fungus (*Beauveria bassiana*), but given the more common use of ITS as a fungal barcoding locus (Seifert 2009) and the probable inefficient amplification of fungal *coxI* when using primers designed for invertebrates, we pooled all fungal hits into one identification and did not consider them further. A small subset of reads was left unassigned by the metaBEAT pipeline. Manual BLAST searches of these sequences through the NCBI website revealed that these were either: (1) Sequences that did not meet the BLAST search criteria (95% similar across at least 90% of the sequence length) due to gaps in database composition; or (2) sequences where a lowest common ancestor could not be assigned due to database error.

**Figure 5.**
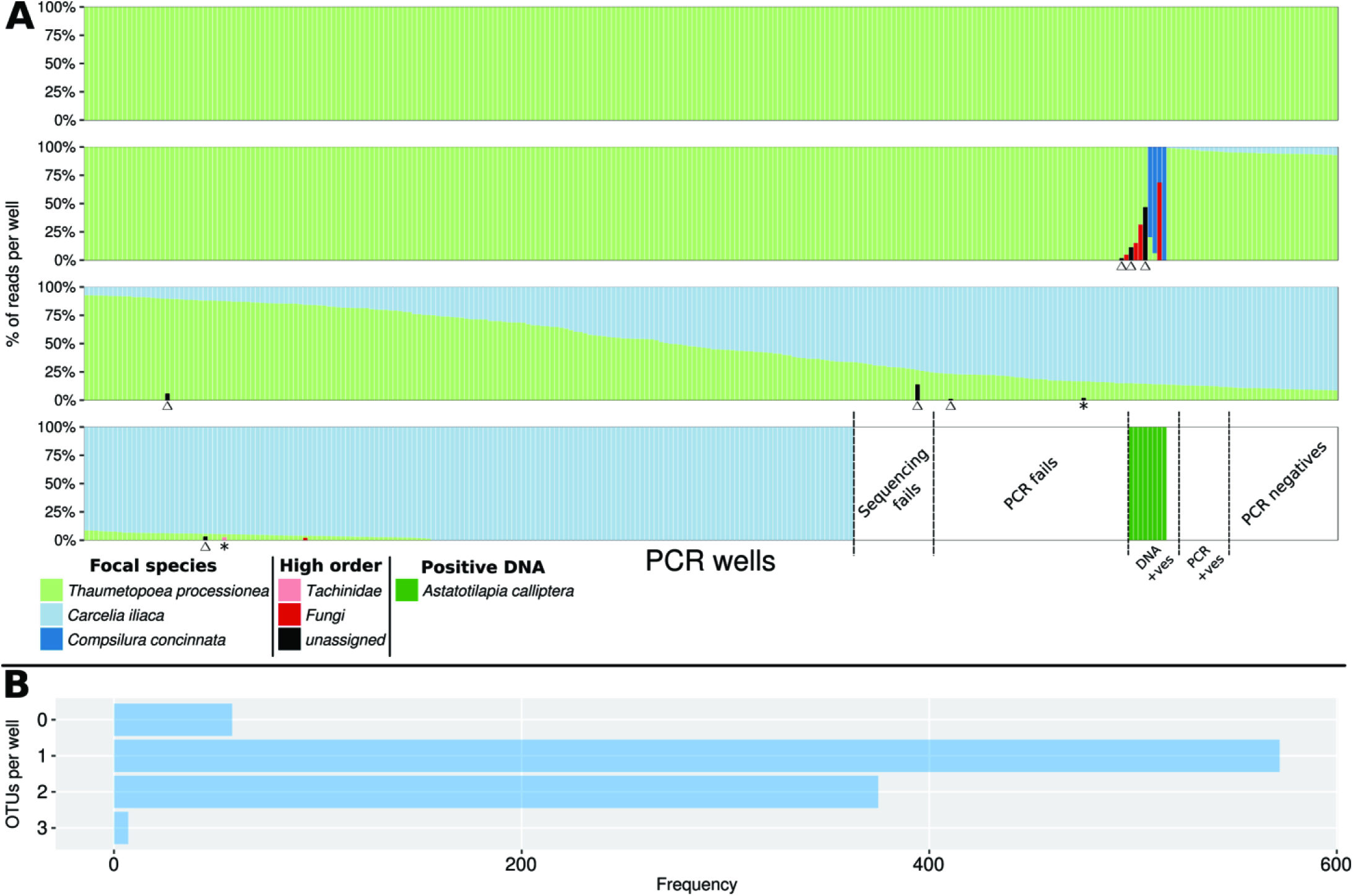
(A) Percentages of reads and taxonomic identifications for all PCR wells. (B) the frequency of different numbers of taxonomic identifications per OPM sample well. Samples in (A) are sorted by descending percentage OPM and increasing percentage *Carcelia iliaca*. Unassigned reads in samples marked with Δ are fungal reads that fall below our BLAST criteria, unassigned reads in the samples marked * are likely to be *Carcelia iliaca* Numts (see main text).

Scenario (1) generally occurs when BLAST identifications are either; all fungal but none are close enough to assign (i.e. probably genuine fungal sequences but from groups poorly represented in Genbank for *coxI* sequences) or dipteran sequences with stop codons in all reading frames suggesting that these are *Carcelia iliaca* NuMts (as defined in Lopez *et al.* 1994). Scenario (2) occurs when the lowest common ancestor algorithm fails because the top 10% of BLAST hits are a mixture of unrelated sequences probably due to the misidentification of sequences in Genbank (e.g. fungal sequences from dipteran specimens labelled as dipteran sequences).

## Discussion

### Evaluating nested metabarcoding for determining Lepidopteran-parasitoid interactions

We tested the ability of a NGS nested metabarcoding design to produce individual-level data for a large number of caterpillar samples (>1000) in a single sequencing run. We achieved a high level of PCR and sequencing success and found an average of 11,000x coverage for each PCR well before sequence filtering, allowing us to adopt a high stringency for sequence quality. The depth of coverage found in our experiment allowed us to distinguish multiple unique sequences in each well, representing the host, parasitoids, and (potentially) any other species interacting with the moths such as parasitic fungi or intracellular parasites. *Thaumetopoea processionea* caterpillars were parasitised by two parasitoid species already known from the literature. *Carcelia iliaca* was found to parasitise almost half of all caterpillars while the other, *Compsilura concinnata*, was only detected in four caterpillars.

In addition to tachinid parasitoids, we detected a range of fungal sequences including the entomopathogenic fungus *Beauveria bassiana.* However, before making assessments of *B. bassiana* infection rates, it would be much better to use fungi specific primers that target the ITS region more commonly used for fungal barcoding (Seifert 2009). Arthropod *coxI* primers are likely to be inefficient at amplifying fungal DNA and the fungal reference libraries are much more complete for ITS. While this would require investment in a set of tagged fungal ITS primers in addition to the general arthropod primers used here, multiple loci may not automatically mean multiple sequencing runs (e.g. Cruaud *et al.* 2017), so the overall cost may not increase considerably, something that is not the case with Sanger approaches. Thus, our approach leads easily to a much more complete understanding of the ecological interactions than standard Sanger barcoding approaches (cf. Wirta *et al.* 2014; Derocles *et al.* 2015). The relative costs of NGS and Sanger sequencing vary with the scale of the experiment. Commercial UK prices for Sanger sequences in both directions are approximately 1/150th the cost of an Illumina MiSeq run at time of writing so for small numbers of individuals and a single barcode locus Sanger sequencing may be much more cost effective. As the quantity of data required increases, however, NGS has the potential to be considerably cheaper, since the costs of a single NGS run are largely fixed, irrespective of how many individuals are included. Although our experiment could have even been performed using Sanger sequencing (through the use of order level primers or cloning), for our experiment we estimate that the costs are at least 1/2 that of the equivalent Sanger experiment even when buying a set of tagged primers sufficient to cover multiple experiments and assuming that no cloning and extra sequencing is required for the Sanger approach (see appendix 4). In reality the cost savings are likely to be much greater.

### Limitations and improvements of the nested metabarcoding approach

Our first attempt at using this method (Supplementary information appendix 1) revealed that it can be highly sensitive to cross contamination between both sample and control wells (see figures S1B, S2B, S4 and S5A; appendix 1 figures are intended for direct comparison with the main text figures). It was suspected that a number of pathways may have contributed to the contamination. First, manual puncturing of caterpillars may be releasing bodily fluids of both host and parasitoid into the air around the DNA extraction plate. We improved this by mechanically lysing caterpillars in closed tubes to contain any aerosols or debris. Second, it was suspected that contaminating aerosols may be moving beneath the commonly used sealing film on a standard 96 well PCR plate during the hottest stages of the PCR cycle. To mitigate this we moved from using 96 well plates to strips of tubes with individual lids and a mineral oil vapour barrier above the PCR mastermix. This improved the quality of the results dramatically allowing us to have much greater confidence that our results are representative of true parasitism rates. In addition to the improvements already implemented here, we would recommend quantification of PCR products using a plate reader and the use of robotic liquid handlers to accurately pool equimolar samples into each pre-library prior to PCR2 as this would likely help control for potential sequencing dropout as possibly seen in our DNA positives.

There is considerable variation in the proportions of reads in each sample attributable to OPM and its parasitoids and it may be the case that this represents true variation in the proportions of each sample composed of OPM or parasitoid tissue but we consider this to be an unreliable approach at present. Some authors have attempted to relate read depth to biomass or numbers of individuals both for PCR based metabarcoding (e.g. Elbrecht & Leese 2015; Thomas *et al.* 2016) and PCR free metabarcoding (e.g. Tang *et al.* 2015). Attempting to measure sample sizes or biomass from read depth presents a number of challenges. First, PCR based approaches can be biased by variation in amplification efficiency across different taxa (for example, variation in primer binding affinities across different taxa or base composition variation affecting enzyme efficiency). PCR free approaches to metabarcoding attempt to circumvent this by removing the PCR step and all the associated biases completely (e.g. Tang *et al.* 2015). In theory, read depth should then correlate with copy number for a given locus, but in reality we have little knowledge, for most species of how sequenceable DNA availability is affected by extraction method and more importantly, how read depth then correlates with biomass or numbers of individuals across different life stages. PCR free metabarcoding is further constrained as much of the read depth which could be used for sequencing additional specimens is used for sequencing additional areas of genome that are not necessary for identification. While much of the variation in proportions of OPM and parasitoid reads in our samples are likely to be attributable to relative proportions of host and parasitoid tissue, we feel that it would be necessary to perform extensive calibration (as in Thomas *et al.* 2014; and Elbrecht & Leese 2015) to make any concrete conclusions surrounding this. Nevertheless, our approach allows us to use presence/absence data across a large number of individual specimens to produce quantitative frequency data that can be analysed with standard statistical tests at the same time as reducing over-sequencing of any single individual.

### OPM, its parasitoids and IPM

Other parasitoids species known to attack OPM in its native range were not detected. Their absence in our data set may be due to our samples being almost exclusively late instar caterpillars, whereas many of the parasitoids recorded in the literature are egg or early instar parasitoids that emerge before nest formation or are pupal parasitoids (Sobczyk 2014). It is not known whether any of the pupal parasitoids of OPM oviposit in late stage larvae then develop after formation of the pupa so it is impossible to say whether recently laid eggs of larval-pupal parasitoids have been missed. It is also always a possibility that there may be false negatives for the detection of very minor components in DNA mixtures, but the improved pooling procedures outlined above would help to mitigate this. In addition, for future screening, it would be useful to explicitly test the detection threshold under laboratory conditions using larvae known to be parasitised and sampled at different time points since parasitism, though this would be extremely difficult given the toxic nature of OPM caterpillars. It is also possible that other UK parasitoid species have yet to colonise OPM. The development of parasitism often exhibits a lag period of some years after the arrival of a new host (e.g. Stone *et al.* 2012 for natural communities; Pocock & Evans 2014 for another example of an invasive lepidopteran species in the UK), and much of the parasitoid community associated with OPM in its native range may simply not be present in the UK, or even if present as a taxonomic species, the local race may be an ecologically adapted cryptic species with different host preferences (e.g. Smith *et al.* 2007). More thorough sampling of all OPM life stages and those of other insect herbivores in the wider forest environment would allow us to assess this. In addition a study examining a series of OPM populations sampled across the native range may provide further insights into potential biocontrol agents for future introduction.

In order to better understand the role of parasitoids in mediating OPM numbers, it is also desirable to consider the place of both OPM and its parasitoids in the broader ecological networks of which they are members, as both direct and indirect interactions with other species in the wider network affect probability of hosts and parasitoids of interest interacting (Hrček & Godfray 2015; Evans *et al.* 2016). This is especially important when considering introducing a new biocontrol agent to an area. By knowing both the alternative hosts of confirmed OPM parasitoids and any previously reported parasitoids, forest managers could design specific planting regimes to enhance parasitoid control of OPM (as suggested in Evans *et al.* 2016). An example of this can be seen with *C. concinnata.* Evidence from the North American use of *C. concinnata* as an introduced biocontrol agent for gypsy and brown-tail moths suggests that this species also has a very wide host range (Strazanac *et al.* 2001; Elkinton & Boettner 2012), and that it is generally ineffective at preventing the spread of the two main target species because of low parasitism levels. This species was also detected at very low levels in UK OPM, but whether these were accidental parasitism events caused by adult females misinterpreting oviposition cues or the first steps in the host range expansion of the UK race of *C. concinnata* is unclear. Understanding both how this will change over time and the competitive effects of other hosts vs OPM for this species is crucial to its evaluation as an OPM biocontrol agent in the UK.

### Nested metabarcoding and ecological networks

Ecological sciences are appreciating more than ever the power of incorporating ecological networks rather than simple species lists into monitoring approaches. The ability to start disentangling species interactions has potential to revolutionise habitat management, habitat restoration, conservation and IPM, but in order to do this there is a need for large well sampled ecological networks (Evans *et al.* 2016). Building such networks requires large sample sizes of individual level rather than community data, and so have previously been little assisted by NGS. Nested metabarcoding can fill this gap and although applied here to parasitised individuals, we anticipate that the sequencing approach demonstrated could be applied in exactly the same way to a range of study systems where it is desirable to sequence numerous samples each containing a restricted number of species, e.g. for detecting pollen on pollinators (current work in prep); identifying recent meals on mouthparts of insect herbivores, or describing interactions between arbuscular mycorrhizal fungi and different plant species. In addition, the networks produced are explicitly linked to sequence data for all the individuals included. This facilitates an evolutionary approach to examining community assembly and for the investigation of broader coevolutionary patterns.

This approach could also be used to process more complex communities in environmental or medical samples, soil mesofauna, bulk insect samples, or any other complex community while still keeping the number of MID tags required at reasonable levels. Should the read number be insufficient for a given experiment, the same samples could be loaded onto a sequencer with higher throughput (e.g. Illumina HiSeq rather than MiSeq) to address this issue, as long as the paired-end nature of the sequences can be maintained. For taxonomic groups that require longer barcodes for accurate identification, emerging technologies such as nanopore sequencing and the PacBio SMRT sequencing may ultimately prove useful.

## Conclusions

Here we demonstrate a highly successful approach to detecting species interactions using a single MiSeq sequencing run. We have shown that a significant proportion of over 1000 OPM caterpillars were parasitised by either *Carcelia iliaca* or *Compsilura concinnata.* The costs are highly favourable compared to undertaking the same study using Sanger based approaches. Scaling this approach would allow for the construction of large, highly-resolved ecological networks of use in a range of applications including conservation and land management, but the sequence based nature of the data generated also allows for the construction of phylogenetically-structured networks that enables many fundamental community dynamics and co-evolutionary questions to be explored. These network and evolutionary based approaches will be of increasing importance as we attempt to quantify functional changes in ecological networks with climate change, habitat modification, and species loss.

## Acknowledgements

We would like to thank Amir Szitenberg for helpful discussion on bioinformatic pipelines and Gillian Jonusas (Royal Parks) for access and logistics in Richmond Park, and Kai Gloyna (State Office for Health and Social Affairs, Rostock, Germany) for providing specimens of *Pales processionea*. We would also like to thank Vasco Elbrecht and two anonymous reviewers for their helpful comments on an earlier version of this manuscript. The project was funded by Forestry Commission England.

## Data Accessibility

To ensure reproducibility of all our analyses we have deposited Jupyter notebooks, R scripts and supplementary material (including tables s1-s4) on Github (<u>Kitson et al. NMB 1.4</u>). An archived version of this release is available on Zenodo (doi: <u>10.5281/zenodo.1066005</u>). Raw sequence data has been submitted to the SRA with accession number <u>PRJNA305686</u>. The metaBEAT pipeline, and other analyses, were run in a Docker container https://hub.docker.com/r/chrishah/metabeat/; v0.97.7 was used for the current study) in order to make our entire analysis environment available for replication if required.

## Author Contributions

DME, DHL were Principal Investigators and designed the project. JJNK collected Croydon specimens in the field, developed protocols and processed specimens in the lab, and undertook analysis and interpretation of the data. CH created metaBEAT and advised on how to process NGS sequence data. NAS facilitated all fieldwork and contributed to project design. RJS collected Richmond park specimens and contributed to specimen processing. All authors contributed to writing and revising the paper.

## References

Binladen J, Gilbert MTP, Bollback JP et al. (2007) The use of coded PCR primers enables high-throughput sequencing of multiple homolog amplification products by 454 parallel sequencing. PloS one, 2, e197.

Boland CRJ (2004) Introduced cane toads *Bufo marinus* are active nest predators and competitors of rainbow bee-eaters *Merops ornatus*: observational and experimental evidence. Biological conservation, 120, 53–62.

Bradshaw CJA, Leroy B, Bellard C et al. (2016) Massive yet grossly underestimated global costs of invasive insects. Nature communications, 7, 12986.

Campbell NR, Harmon SA, Narum SR (2015) Genotyping-in-Thousands by sequencing (GT-seq): A cost effective SNP genotyping method based on custom amplicon sequencing. Molecular ecology resources, 15, 855–867.

Cruaud P, Rasplus J-Y, Rodriguez LJ, Cruaud A (2017) High-throughput sequencing of multiple amplicons for barcoding and integrative taxonomy. Scientific reports, 7, 41948.

Day WH (1994) Estimating Mortality Caused by Parasites and Diseases of Insects: Comparisons of the Dissection and Rearing Methods. Environmental entomology, 23, 543–550.

Dejean T, Valentini A, Miquel C et al. (2012) Improved detection of an alien invasive species through environmental DNA barcoding: the example of the American bullfrog *Lithobates catesbeianus*. The Journal of applied ecology, 49, 953–959.

Derocles SAP, Evans DM, Nichols PC, Evans SA, Lunt DH (2015) Determining plant - leaf miner - parasitoid interactions: a DNA barcoding approach. PloS one, 10, e0117872.

Elbrecht V, Leese F (2015) Can DNA-Based Ecosystem Assessments Quantify Species Abundance? Testing Primer Bias and Biomass--Sequence Relationships with an Innovative Metabarcoding Protocol. PloS one, 10, e0130324.

Elkinton JS, Boettner GH (2012) Benefits and harm caused by the introduced generalist tachinid, Compsilura concinnata, in North America. Biocontrol, 57, 277–288.

Evans DM, Kitson JJN, Lunt DH, Straw NA, Pocock MJO (2016) Merging DNA metabarcoding and ecological network analysis to understand and build resilient terrestrial ecosystems. Functional ecology, 30, 1904–1916.

Fadrosh DW, Ma B, Gajer P et al. (2014) An improved dual-indexing approach for multiplexed 16S rRNA gene sequencing on the Illumina MiSeq platform. Microbiome, 2, 6.

Forest Research (2017) Oak Processionary Moth Programme: Operational Report 2016. OPM Programme Board.

Giguet-Covex C, Pansu J, Arnaud F et al. (2014) Long livestock farming history and human landscape shaping revealed by lake sediment DNA. Nature communications, 5, 3211.

Groenen F, Meurisse N (2012) Historical distribution of the oak processionary moth Thaumetopoea processionea in Europe suggests recolonization instead of expansion. Agricultural and forest entomology, 14, 147–155.

Handley L-JL, Estoup A, Evans DM et al. (2011) Ecological genetics of invasive alien species. Biocontrol, 56, 409–428.

Hesketh H, Roy HE, Eilenberg J, Pell JK, Hails RS (2010) Challenges in modelling complexity of fungal entomopathogens in semi-natural populations of insects. Biocontrol, 55, 55–73.

Hrček J, Godfray HCJ (2015) What do molecular methods bring to host-parasitoid food webs? Trends in parasitology, 31, 30–35.

Hyslop EJ (1980) Stomach contents analysis–a review of methods and their application. Journal of fish biology.

Illumina (2011) Preparing 16S Ribosomal RNA Gene Amplicons for the Illumina MiSeq System. Illumina technical note.

Kaartinen R, Stone GN, Hearn J, Lohse K, Roslin T (2010) Revealing secret liaisons: DNA barcoding changes our understanding of food webs. Ecological entomology, 35, 623–638.

Lamy M, Pastureaud MH, Novak F et al. (1986) Thaumetopoein: an urticating protein from the hairs and integument of the pine processionary caterpillar (*Thaumetopoea pityocampa* Schiff., Lepidoptera, Thaumetopoeidae). Toxicon: official journal of the International Society on Toxinology, 24, 347–356.

Lefort M-C, Wratten SD, Cusumano A, Varennes Y-D, Boyer S (2017) Disentangling higher trophic level interactions in the cabbage aphid food web using high-throughput DNA sequencing. Metabarcoding and Metagenomics, 1, e13709.

Leray M, Yang JY, Meyer CP et al. (2013) A new versatile primer set targeting a short fragment of the mitochondrial COI region for metabarcoding metazoan diversity: application for characterizing coral reef fish gut contents. Frontiers in zoology, 10, 34.

Lopezaraiza–Mikel ME, Hayes RB, Whalley MR, Memmott J (2007) The impact of an alien plant on a native plant–pollinator network: an experimental approach. Ecology letters, 10, 539–550.

Lopez JV, Yuhki N, Masuda R, Modi W, O’Brien SJ (1994) Numt, a recent transfer and tandem amplification of mitochondrial DNA to the nuclear genome of the domestic cat. Journal of molecular evolution, 39, 174–190.

Maglianesi MA, Blüthgen N, Böhning-Gaese K, Schleuning M (2014) Morphological traits determine specialization and resource use in plant–hummingbird networks in the neotropics. Ecology, 95, 3325–3334.

Mindlin MJ, le Polain de Waroux O, Case S, Walsh B (2012) The arrival of oak processionary moth, a novel cause of itchy dermatitis, in the UK: experience, lessons and recommendations. Public health, 126, 778–781.

Otte D, Joern A (1976) On Feeding Patterns in Desert Grasshoppers and the Evolution of Specialized Diets. Proceedings of the Academy of Natural Sciences of Philadelphia, 128, 89–126.

Peck HL, Pringle HE, Marshall HH, Owens IPF, Lord AM (2014) Experimental evidence of impacts of an invasive parakeet on foraging behavior of native birds. Behavioral ecology: official journal of the International Society for Behavioral Ecology, 25, 582–590.

Pocock MJO, Evans DM (2014) The success of the horse-chestnut leaf-miner, Cameraria ohridella, in the UK revealed with hypothesis-led citizen science. PloS one, 9, e86226.

Pocock MJO, Evans DM, Memmott J (2012) The robustness and restoration of a network of ecological networks. Science, 335, 973–977.

Rognes T, Flouri T, Nichols B, Quince C, Mahé F (2016) VSEARCH: a versatile open source tool for metagenomics. PeerJ, 4, e2584.

Roques A (2010) Taxonomy, time and geographic patterns. Chapter 2. In: Alien terrestrial arthropods of Europe (eds Roques A, Kenis M, Lees D, et al.), p. 11. Pensoft Publishers.

Roques A (2014) Processionary Moths and Climate Change: An Update. Springer.

Roques A, Auger-Rozenberg M-A, Blackburn TM et al. (2016) Temporal and interspecific variation in rates of spread for insect species invading Europe during the last 200 years. Biological invasions, 18, 907–920.

Roy HE, Hails RS, Hesketh H, Roy DB, Pell JK (2009) Beyond biological control: non-pest insects and their pathogens in a changing world. Insect conservation and diversity / Royal Entomological Society of London, 2, 65–72.

Roy HE, Preston CD, Harrower CA et al. (2014) GB Non-native Species Information Portal: documenting the arrival of non-native species in Britain. Biological invasions, 16, 2495–2505.

Sands RJ, Jonusas G, Straw NA, Kitson JJN, Raper CM (2015) *Carcelia Iliaca* (Diptera: Tachinidae), a specific parasitoid of the Oak Processionary Moth (Lepidoptera: Thaumetopoeidae), new to Great Britain. British Journal of Entomology and Natural History, 28, 225–227.

Schnell IB, Bohmann K, Gilbert MTP (2015) Tag jumps illuminated - reducing sequence-to-sample misidentifications in metabarcoding studies. Molecular ecology resources, 15, 1289–1303.

Schooler SS, De Barro P, Ives AR (2011) The potential for hyperparasitism to compromise biological control: Why don’t hyperparasitoids drive their primary parasitoid hosts extinct? Biological control: theory and applications in pest management, 58, 167–173.

Seifert KA (2009) Progress towards DNA barcoding of fungi. Molecular ecology resources, 9 Suppl s1, 83–89.

Shokralla S, Porter TM, Gibson JF et al. (2015) Massively parallel multiplex DNA sequencing for specimen identification using an Illumina MiSeq platform. Scientific reports, 5, 9687.

Šigut M, Kostovčík M, Šigutová H et al. (2017) Performance of DNA metabarcoding, standard barcoding, and morphological approach in the identification of host-parasitoid interactions. PloS one, 12, e0187803.

Smith MA, Rodriguez JJ, Whitfield JB et al. (2008) Extreme diversity of tropical parasitoid wasps exposed by iterative integration of natural history, DNA barcoding, morphology, and collections. Proceedings of the National Academy of Sciences of the United States of America, 105, 12359–12364.

Smith MA, Wood DM, Janzen DH, Hallwachs W, Hebert PDN (2007) DNA barcodes affirm that 16 species of apparently generalist tropical parasitoid flies (Diptera, Tachinidae) are not all generalists. Proceedings of the National Academy of Sciences of the United States of America, 104, 4967–4972.

Sobczyk T (2014) Der Eichenprozessionsspinner in Deutschland. Bundesamt für Naturschutz.

Stigter H, Geraedts W, Spijkers H (1997) *Thaumetopoea processionea* in the Netherlands: present status and management perspectives (Lepidoptera: Notodontidae). Proceedings of the Section Experimental and Applied Entomology of the Netherlands Entomological Society, 8, 3–16.

Stone GN, Lohse K, Nicholls JA et al. (2012) Reconstructing Community Assembly in Time and Space Reveals Enemy Escape in a Western Palearctic Insect Community. Current biology: CB, 22, 532–537.

Strazanac JS, Plaugher CD, Petrice TR, Butler L (2001) New Tachinidae (Diptera) host records of eastern North American forest canopy Lepidoptera: baseline data in a Bacillus thuriengiensis variety kurstaki nontarget study. Journal of economic entomology, 94, 1128–1134.

Taberlet P, Coissac E, Pompanon F, Brochmann C, Willerslev E (2012) Towards next-generation biodiversity assessment using DNA metabarcoding. Molecular ecology, 21, 2045–2050.

Tang M, Hardman CJ, Ji Y et al. (2015) High-throughput monitoring of wild bee diversity and abundance via mitogenomics. Methods in ecology and evolution / British Ecological Society, 6, 1034–1043.

Thomas AC, Deagle BE, Eveson JP, Harsch CH, Trites AW (2016) Quantitative DNA metabarcoding: improved estimates of species proportional biomass using correction factors derived from control material. Molecular ecology resources, 16, 714–726.

Thomas AC, Jarman SN, Haman KH, Trites AW, Deagle BE (2014) Improving accuracy of DNA diet estimates using food tissue control materials and an evaluation of proxies for digestion bias. Molecular ecology, 23, 3706–3718.

Traugott M, Bell JR, Broad GR et al. (2008) Endoparasitism in cereal aphids: molecular analysis of a whole parasitoid community. Molecular ecology, 17, 3928–3938.

Truett GE, Heeger P, Mynatt RL et al. (2000) Preparation of PCR-quality mouse genomic DNA with hot sodium hydroxide and tris (HotSHOT). BioTechniques, 29, 52, 54.

Vazquez-Prokopec GM, Chaves LF, Ritchie SA, Davis J, Kitron U (2010) Unforeseen costs of cutting mosquito surveillance budgets. PLoS neglected tropical diseases, 4, e858.

Wagenhoff E, Veit H (2011) Five Years of Continuous *Thaumetopoea processionea* Monitoring: Tracing Population Dynamics in an Arable Landscape of South-Western Germany. Gesunde Pflanzen, 63, 51–61.

Wirta HK, Hebert PDN, Kaartinen R et al. (2014) Complementary molecular information changes our perception of food web structure. Proceedings of the National Academy of Sciences of the United States of America, 111, 1885–1890.

Yu DW, Ji Y, Emerson BC et al. (2012) Biodiversity soup: metabarcoding of arthropods for rapid biodiversity assessment and biomonitoring. Methods in ecology and evolution / British Ecological Society, 3, 613–623.

Zwakhals CJ (2005) *Pimpla processioneae* and *P. rufipes*: specialist versus generalist (Hymenoptera: Ichneumonidae, Pimplinae). Entomologische Berichten, 65, 14–16.

